# The Cytochrome b m.14849T>C (S35P) Variant Induces Structural and Dynamic Alterations in the Heme bL Microenvironment in Multisystem Disease

**DOI:** 10.64898/2026.02.27.708559

**Authors:** Ekrem Yasar, Abdussamed Yasin Demir, Segun Dogru

**Affiliations:** Department of Biophysics, Erzincan Binali Yildirim University, Erzincan, Turkey; Department of Medical Genetics, Erzincan Binali Yildirim University, Erzincan, Turkey; Department of Medical Biochemistry, Akdeniz University Medical School, Antalya, Turkey

**Keywords:** MT-CYB, Variant of Uncertain Significance, Heme bL Microenvironment, Molecular Dynamics Simulation, Mitochondrial Complex III

## Abstract

Mitochondrial Complex III dysfunction is frequently associated with pathogenic variants in the MT-CYB gene, yet the functional consequences of many missense substitutions remain unresolved because they are classified as variants of uncertain significance (VUS). One such variant, m.14849T>C (p.Ser35Pro), has been reported in patients with multisystem mitochondrial phenotypes, including septo-optic dysplasia, cardiomyopathy, and exercise intolerance, although its structural impact on Cytochrome b function remains unclear.

In this study, we employed 300 ns all-atom molecular dynamics simulations to assess structural and energetic consequences of the S35P substitution in the Cytochrome b subunit of human mitochondrial Complex III. The S35P variant did not induce global destabilization of the protein scaffold but instead promoted localized perturbations within the heme bL microenvironment. The mutation was associated with loss of a heme-proximal hydrogen-bonding network involving Ser35 and a decrease in electrostatic interaction energy between the protein matrix and the heme bL cofactor. Radial distribution function analysis further supported loosening of local packing around the prosthetic group.

Consistent with these local changes, dynamics analyses indicated increased flexibility in distal transmembrane helices that form the heme-pocket scaffold and greater variability in the inter-heme Fe(bL)–Fe(bH) distance. Together, our findings suggest that S35P may exert functional effects by reorganizing the heme bL microenvironment rather than by inducing large-scale structural destabilization, underscoring the value of structure- and dynamics-based evaluation for mitochondrial VUS and suggesting a plausible mechanistic link to the pathophysiology of multisystem mitochondrial diseases.

## 1 INTRODUCTION

Mitochondrial oxidative phosphorylation (OXPHOS) is the primary source of ATP production in eukaryotic cells and depends on the coordinated function of five multi-subunit respiratory chain complexes embedded in the inner mitochondrial membrane [1-3]. Among these, Complex III (cytochrome bc1 complex) plays a pivotal role in electron transfer by catalyzing the oxidation of ubiquinol and reduction of cytochrome c through the Q-cycle mechanism [4-6]. This process critically relies on the structural integrity and dynamic coordination of redox-active cofactors, including heme bL and heme bH, which are embedded within the mitochondrial cytochrome b (MT-CYB) subunit [7-9].

The MT-CYB gene, encoded by mitochondrial DNA (mtDNA), constitutes the only mitochondrially encoded core catalytic component of Complex III and is essential for maintaining efficient electron transfer and proton gradient formation across the inner mitochondrial membrane [10; 11]. Pathogenic variants in MT-CYB have been associated with a wide spectrum of mitochondrial disorders, including exercise intolerance, cardiomyopathy, multisystem metabolic syndromes, and neurodegenerative conditions [12-17]. However, despite the increasing number of clinically reported MT-CYB variants, the molecular mechanisms linking genotype to impaired electron transport and mitochondrial dysfunction remain poorly understood [18-20].

One such variant is the m.14849T>C transition, resulting in a serine-to-proline substitution at position 35 (p.Ser35Pro; S35P) in the MT-CYB protein. This variant has been reported in association with multisystem mitochondrial phenotypes including exercise intolerance and cardiomyopathy, and has also been described in patients presenting with septo-optic dysplasia and isolated Complex III deficiency [13; 14; 21]. Nevertheless, due to the absence of functional or mechanistic characterization, the S35P substitution is currently classified as a variant of uncertain significance (VUS) in mitochondrial variant databases such as MITOMAP and ClinVar [22; 23]. Interpreting the functional consequences of such VUS remains a major challenge in mitochondrial genetics, particularly for variants located within structurally or dynamically sensitive regions of respiratory chain proteins [24; 25].

Recent studies have highlighted that disease-associated mutations in mitochondrial proteins may not necessarily induce global structural destabilization but instead impair function by perturbing conformational dynamics, cofactor coordination, or local electrostatic environments required for efficient redox activity [26; 27]. In the context of MT-CYB, the spatial arrangement and dynamic adaptability of heme b cofactors are critical determinants of electron transfer efficiency and are sensitive to subtle alterations in local secondary structure and hydrogen-bonding networks [28; 29]. Therefore, investigating mutation-induced changes in the structural and energetic properties of MT-CYB may provide valuable mechanistic insight into the functional impact of clinically reported variants.

In this study, we aimed to elucidate the molecular consequences of the MT-CYB m.14849T>C (S35P) variant using comparative all-atom molecular dynamics simulations of wild-type and mutant proteins embedded in a membrane environment. By integrating structural, dynamic, and energetic analyses, we sought to provide a molecular-level functional interpretation of this clinically reported MT-CYB variant and contribute to bridging the gap between genetic variation and mitochondrial electron transport dysfunction [21].

## 2. MATERIALS AND METHODS

### 2.1. Structural Model Preparation and Mutagenesis

The initial coordinates for the human mitochondrial cytochrome b (MT-CYB) were retrieved from the high-resolution cryo-electron microscopy structure of the human kidney respiratory complex III (PDB ID: 9CG3) [30]. The protein sequence corresponds to UniProt ID: P00156 (CYB_HUMAN). Missing terminal residues (Met1 and Leu380) were modeled using the CHARMM-GUI PDB Reader & Manipulator tool to ensure a complete polypeptide chain. The p.Ser35Pro (S35P) mutation was introduced at the structural level using the CHARMM-GUI program [31]. Both the wild-type (WT) and mutant structures were processed to ensure correct protonation states at physiological pH.

### 2.2. Membrane System Assembly and Solvation

To simulate the protein in a biologically relevant environment, both systems were embedded into a complex asymmetric lipid bilayer using the CHARMM-GUI Membrane Builder [32]. The membrane composition was designed to approximate the inner mitochondrial membrane (IMM) environment and consisted of a POPC:POPE:TOCL2 mixture in a 45:35:20 molar ratio. POPC (1-palmitoyl-2-oleoyl-sn-glycero-3-phosphocholine) and POPE (1-palmitoyl-2-oleoyl-sn-glycero-3-phosphoethanolamine) were included to represent the major zwitterionic phospholipids, while cardiolipin (tetraoleoyl cardiolipin, TOCL2), a signature lipid essential for respiratory complex stability, was incorporated to capture key structural features of the IMM. The systems were solvated with ∼22,500 TIP3P water molecules and neutralized with 0.15 M KCl ions. The final simulation boxes (96 × 96 × 110 Å^3^) comprised approximately 106,000 atoms, providing a structurally and compositionally representative model of the mitochondrial inner membrane microenvironment.

### 2.3. Molecular Dynamics Simulation

All simulations were performed using the GROMACS 2024.4 software package [33] with the CHARMM36m all-atom force field [34]. The initial systems underwent energy minimization using the steepest descent algorithm, followed by multi-step equilibration in NVT and NPT ensembles with harmonic restraints on the protein and lipids, following the standard CHARMM-GUI protocol. Production runs were executed for 300 ns for each system under periodic boundary conditions. Temperature and pressure were maintained at 310 K and 1 bar using the Nosé-Hoover thermostat [35] and Parrinello-Rahman barostat [36], respectively. Long-range electrostatic interactions were calculated using the Particle Mesh Ewald (PME) method [37] and covalent bond constraints were applied using the LINCS algorithm [38]. Acceleration was achieved using NVIDIA A4000 GPU units.

### 2.4. Trajectory and Conformational Analysis

Post-simulation analyses were conducted using standard GROMACS tools and custom scripts [39]. Structural stability and flexibility were assessed via Root Mean Square Deviation (RMSD), Root Mean Square Fluctuation (RMSF), and Radius of Gyration (Rg). The solvent-accessible surface area (SASA) and radial distribution function (RDF) were calculated to monitor the hydration patterns [40]. To quantify the mutation induced bending of Helix-A, kink analyses [41] were performed using the gmx helix tool. Essential dynamics were explored through Principal Component Analysis (PCA) [42]. The covariance matrix was constructed using the gmx_covar tool, and the first two principal components (PC1 and PC2) were extracted via gmx_anaeig to visualize the conformational clusters. The conformational free energy landscape of WT and S35P MT-CYB systems was constructed using RMSD and Rg as reaction coordinates based on the Boltzmann distribution principle [43]. The relative Gibbs free energy (ΔG) was calculated from the probability distribution P(x) of conformational states according to:

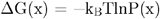

where k_B_ is the Boltzmann constant, T is the temperature (310 K), and P(x) represents the probability distribution of the selected reaction coordinates.

Structural visualization, residue–cofactor interaction analysis, hydrogen-bond inspection, and comparative structural representations of the wild-type and S35P mutant systems were performed using equilibrated molecular dynamics trajectory snapshots in The PyMOL Molecular Graphics System, Version 2.5.2 (Schrödinger, LLC) [44].

### 2.5. Interaction and Energetic Analysis

To quantify the thermodynamic stability of the protein–cofactor interaction, we employed the Molecular Mechanic/Poisson Boltzmann Surface Area (MM-PBSA) method [45] via the gmx_MMPBSA package [46]. Binding free energies were computed based on the thermodynamic cycle summarized by the following equation:

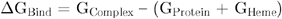

where G_Complex_ represents the total free energy of the MT-CYB–Heme complex, and G_Protein_ and G_Heme_ denote the total free energies of the receptor and the Heme prosthetic group, respectively. The total binding energy was decomposed into four key components: (i) Van der Waals, (ii) electrostatic interactions, (iii) polar solvation, and (iv) non-polar solvation energies. A dielectric constant of 80 (solvent) and a grid spacing of 0.5 Å were applied consistently for accurate electrostatic calculations.

Hydrogen bonding patterns between Residue 35 and the surrounding protein environment were quantified using gmx_hbnum and Visual Molecular Dynamics (VMD) version 2.0 program [47] hydrogen bond occupancy analysis. Inter-atomic distances between Residue 35 and the Heme bL prosthetic group, as well as the Fe−Fe distance between Heme cofactors (Heme bL and Heme bH), were monitored using gmx_distance. Secondary structure transitions and molecular movies were generated using the VMD Timeline plugin [48] and rendering tools.

### 2.6. Sequence Alignment and Clinical Data Curation

Pairwise sequence alignment of MT-CYB was performed using the BLAST algorithm [49] to assess the evolutionary conservation of the Ser35 residue. Clinical data associated with the m.14849T>C variant were curated from the MITOMAP [22], ClinVar [23], MedGen [50], ClinGen [51], OMIM [52] and gnomAD [53] databases. Reported phenotypes and pathogenic classifications were summarized in Table S1 to provide clinical context for the computational findings.

## 3 RESULTS

### 3.1. Structural Framework and Evolutionary Context of the S35P Variant

To establish a baseline for our atomistic investigation, we first confirmed that the wild-type (WT) and p.Ser35Pro (S35P) mutant sequences exhibit near-complete identity (99.7%), isolating the S35P substitution as the sole driver of the observed structural and dynamical differences (Fig. S1). While clinically cataloged as a variant of uncertain significance (VUS), the strategic positioning of Ser35 within the heme-proximal Helix-A suggests its potential role as a structural anchor. Our comparative modeling aims to decipher how this single-point transition from a polar, hydrogen-bonding serine to a proline residue with restricted backbone conformational flexibility reshapes the local microenvironment of the Cytochrome b subunit (Table S1).

Structural inspection of the Cytochrome b heme bL pocket revealed that residue 35 is positioned in close spatial proximity to the cofactor-binding environment. Comparative visualization of WT and S35P mutant structures demonstrated that the substitution occurs within a helix directly contributing to the heme-proximal scaffold, without inducing large-scale structural deviation in the initial conformation (Fig. 1). Spatial mapping further indicates that residue 35 is positioned within Helix-A (residues 28-53) in proximity to the heme bL pocket, while distal helices forming the opposing scaffold (residues 90–120) are dynamically affected in the mutant system (Fig. 1).

**Fig. 1.**
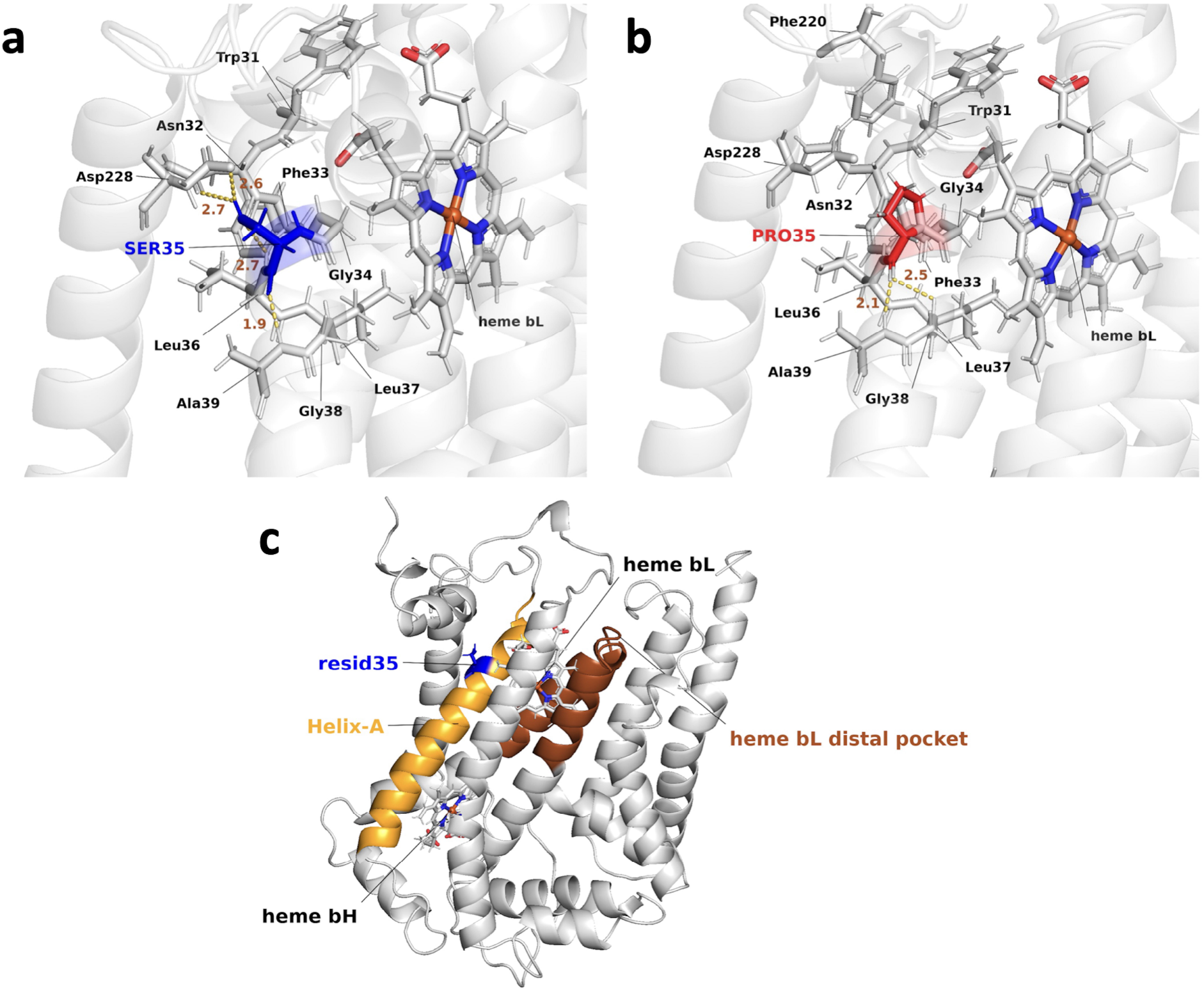
Structural context of the p.Ser35Pro substitution within the heme bL region of Cytochrome b. (a) Local interaction network of Ser35 in the wild-type (WT) structure showing hydrogen-bonding contacts with surrounding residues, including Asp228 and Asn32, in proximity to the heme bL cofactor. (b) Corresponding view of the S35P mutant (Pro35) illustrating disruption of the local interaction network due to substitution of serine with proline, resulting in reduced hydrogen-bonding capacity within the heme bL–proximal environment. (c) Side-view structural representation of Cytochrome b highlighting the spatial localization of residue 35 within Helix-A (residues 28–53) and its relationship to the heme bL distal pocket (residues 90–120). The heme bL and heme bH cofactors are shown for structural reference.

### 3.2. Paradoxical Stability: Global Rigidification vs. Local Helical Distortion

Analysis of backbone RMSD trajectories revealed distinct divergent behaviors between the two systems (Fig. 2). The WT system exhibited characteristic conformational plasticity, traversing a transition state after ∼200 ns to sample a broader conformational space, with RMSD values reaching up to ∼4.5 Å. In contrast, the S35P mutant was characterized by a restricted RMSD profile, fluctuating within a narrow ∼3.0–3.8 Å band. Rather than indicating superior thermodynamic stability, this reduced deviation is consistent with a constrained conformational ensemble.

**Fig. 2.**
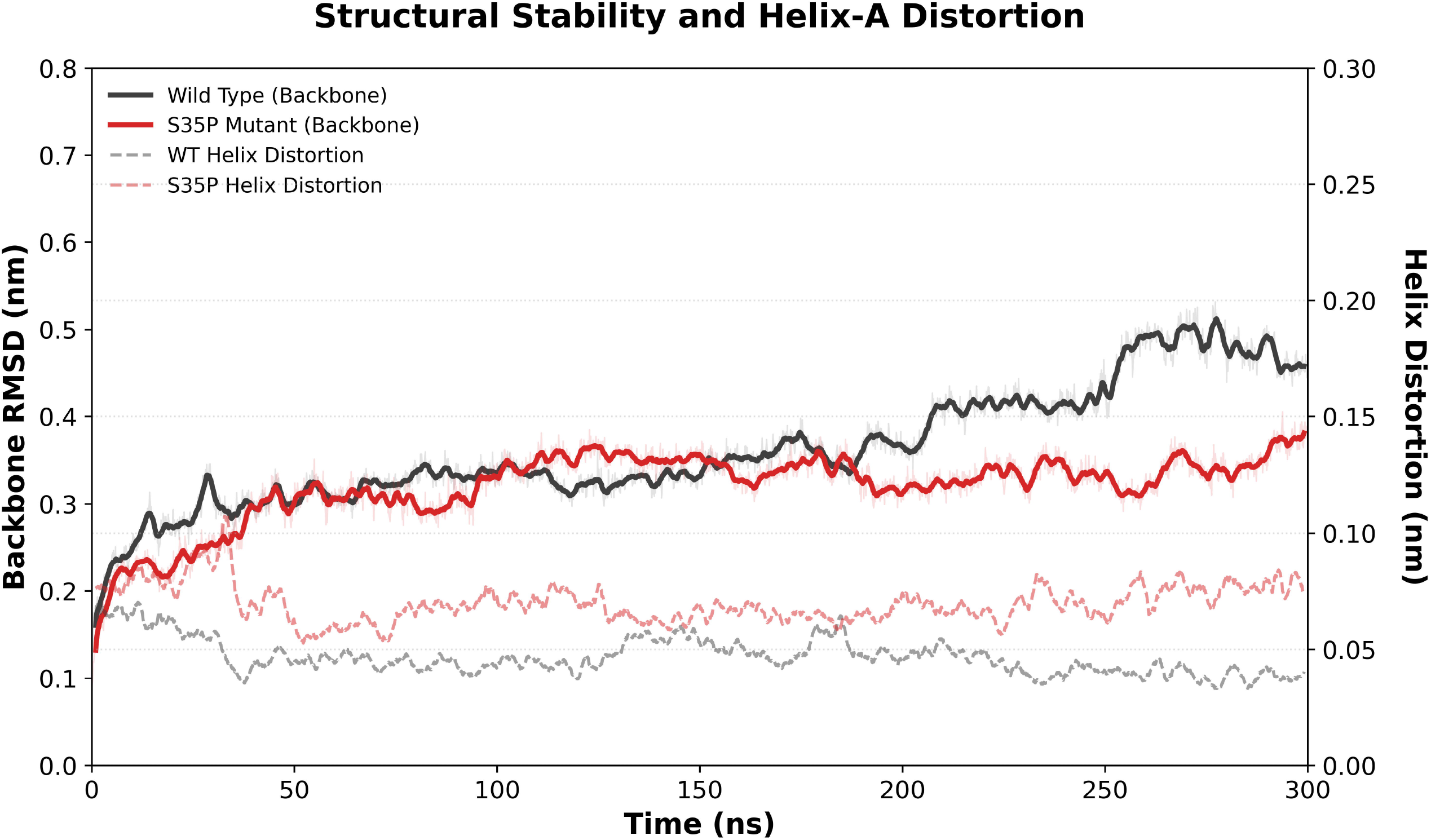
Structural stability and Helix-A distortion of MT-CYB during 300-ns simulations. Time evolution of backbone RMSD for WT (black) and S35P mutant (red) MT-CYB. Helix-A distortion over time (right y-axis; dashed lines) for WT (gray dashed) and S35P (red dashed).

This global rigidification is paradoxically coupled with an increase in Helix-A distortion in the mutant system (Fig. 2). The proline residue induces a localized kink in the helical geometry, elevating the distortion metric to approximately ∼0.8 Å. Importantly, global metrics such as the Radius of Gyration (Rg) and SASA support that while the protein maintains its overall fold without global structural collapse (Fig. S2, Fig. S3), the S35P variant may compromise the local helical persistence near the cofactor-binding environment.

### 3.3. Allosteric Coupling and Dynamic Redistribution

The impact of the S35P mutation is not confined to the mutation site but propagates through the transmembrane bundle. RMSF profiles indicated that while the average mobility of Helix-A remains comparable, the mutant induces elevated fluctuations in a distal segment (residues 90–120) that forms the opposing structural scaffold of the heme bL pocket (Fig. 3). Secondary structure timeline analysis further revealed that this distal destabilization is rooted in increased helix-to-coil transitions (Fig. S4). This long-range dynamic coupling suggests that the S35P substitution disrupts the synchronized conformational dynamics of the heme bL pocket, which could influence heme-proximal structural adaptability relevant to the Q-cycle, the electron transfer process within Complex III that occurs between the heme bL and heme bH cofactors.

**Fig. 3.**
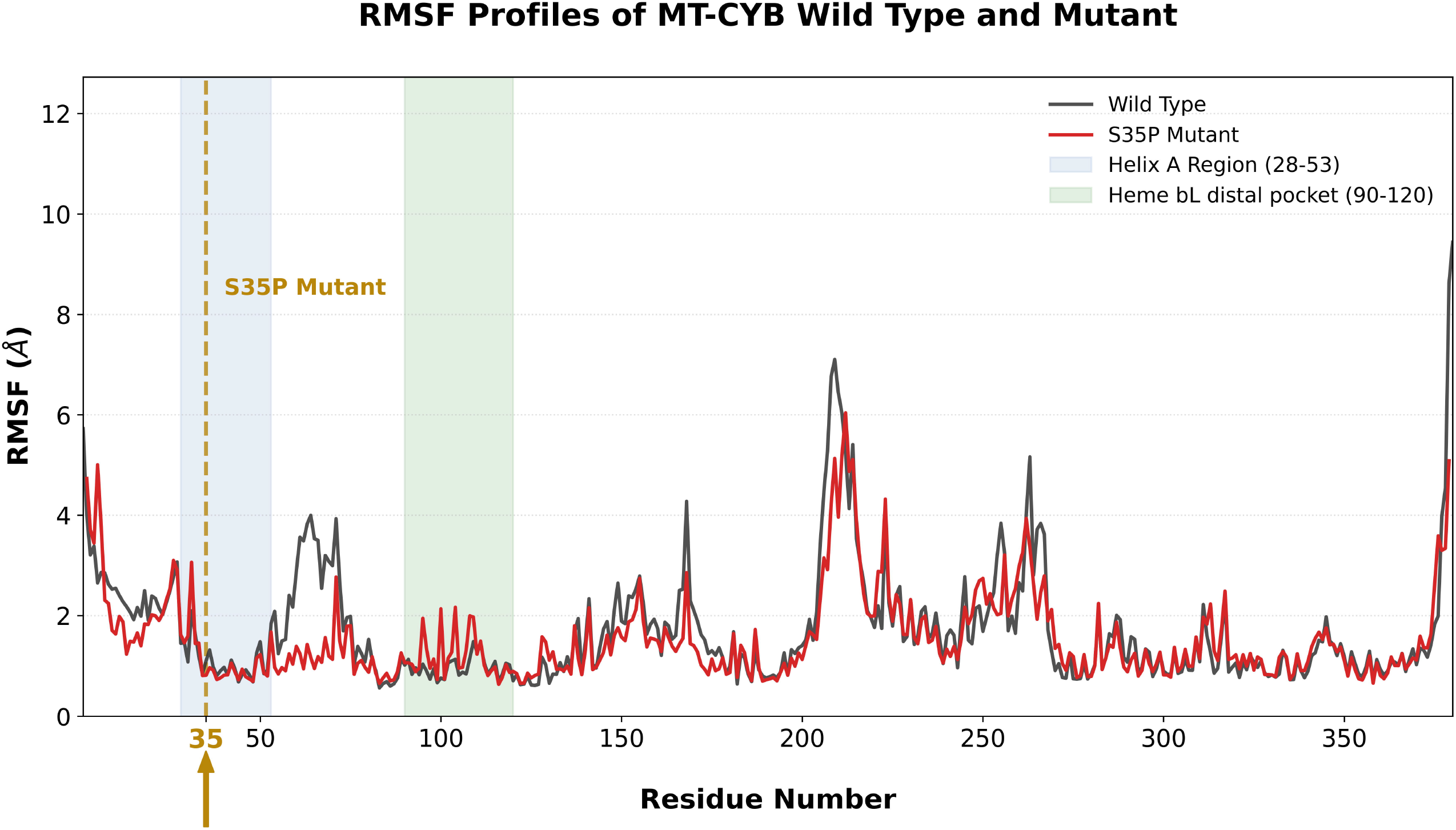
Residue-wise RMSF profiles highlight mutation-induced dynamical redistribution around the heme bL environment. Backbone RMSF (Å) calculated over 300-ns simulations for WT (black) and S35P mutant (red) MT-CYB. The Helix-A region is indicated (blue shaded area) and residue 35 is marked by a dashed vertical line.

### 3.4. Deconstruction of the Ser35 Interaction Hub

The primary molecular defect of the S35P variant was identified as a marked disruption of a critical interaction hub. In the WT system, Ser35 acts as a polar tether, forming a robust hydrogen-bonding network with an average of ∼1.90 bonds, primarily involving Asp228 (∼45% occupancy) and Asn32 (Fig. 4, Fig. S5). The S35P mutation, by introducing a proline residue devoid of side-chain donor capacity, substantially reduces these electrostatic anchors (occupancy <0.1%). This interaction loss is consistent with the subsequent energetic and geometric perturbations observed in the heme bL environment.

**Fig. 4.**
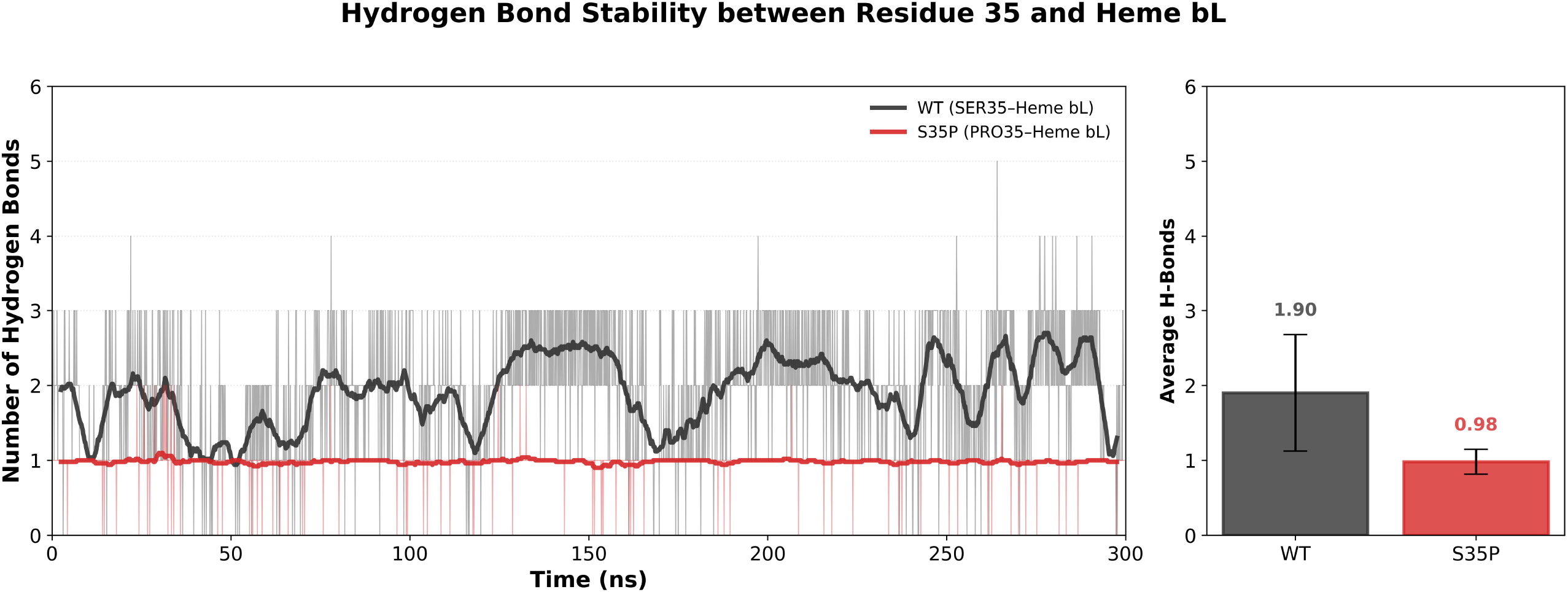
Hydrogen bond stability between residue 35 and the heme bL cofactor. (a) Time evolution of hydrogen bonds formed between residue 35 and heme bL for WT (Ser35; black) and S35P mutant (Pro35; red) over 300 ns. (b) Average number of hydrogen bonds (mean ± SD) formed between residue 35 and heme bL in WT and S35P systems.

### 3.5. Conformational Confinement and Energy Landscape Restriction

To visualize the global impact on the protein’s functional adaptability, we projected the dynamics onto the Principal Component (PC) space and Free Energy Landscapes (FEL). PCA of backbone dynamics revealed that while WT conformations occupied a broad and continuous PC1–PC2 distribution, the S35P mutant exhibited discrete and clustered populations (Fig. 5).

**Fig. 5.**
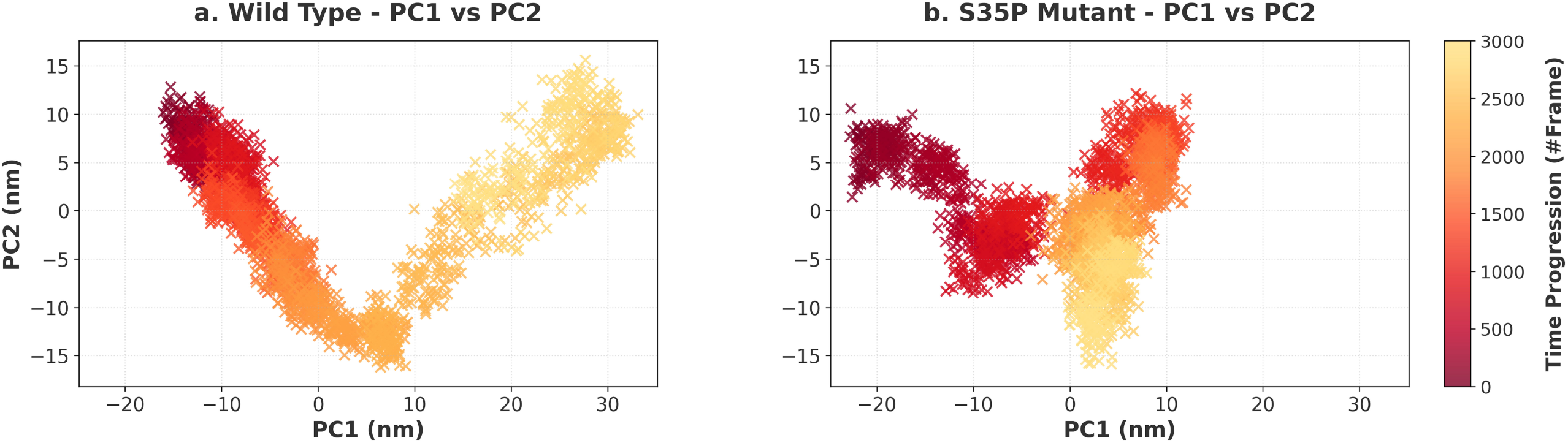
Principal component analysis of WT and S35P MT-CYB. Projection of trajectory frames onto the first two principal components (PC1 and PC2) for WT (left) and S35P mutant (right) systems. Points are colored according to simulation time progression.

FEL analysis further supported this conformational confinement, with the WT system sampling a wide, continuous energy basin, whereas the S35P variant displayed a confined and sharply bounded minimum-energy region (Fig. 6). This conformational confinement is consistent with a reduced ability of the mutant system to sample diverse configurations relevant to structural adaptability within the Complex III framework.

**Fig. 6.**
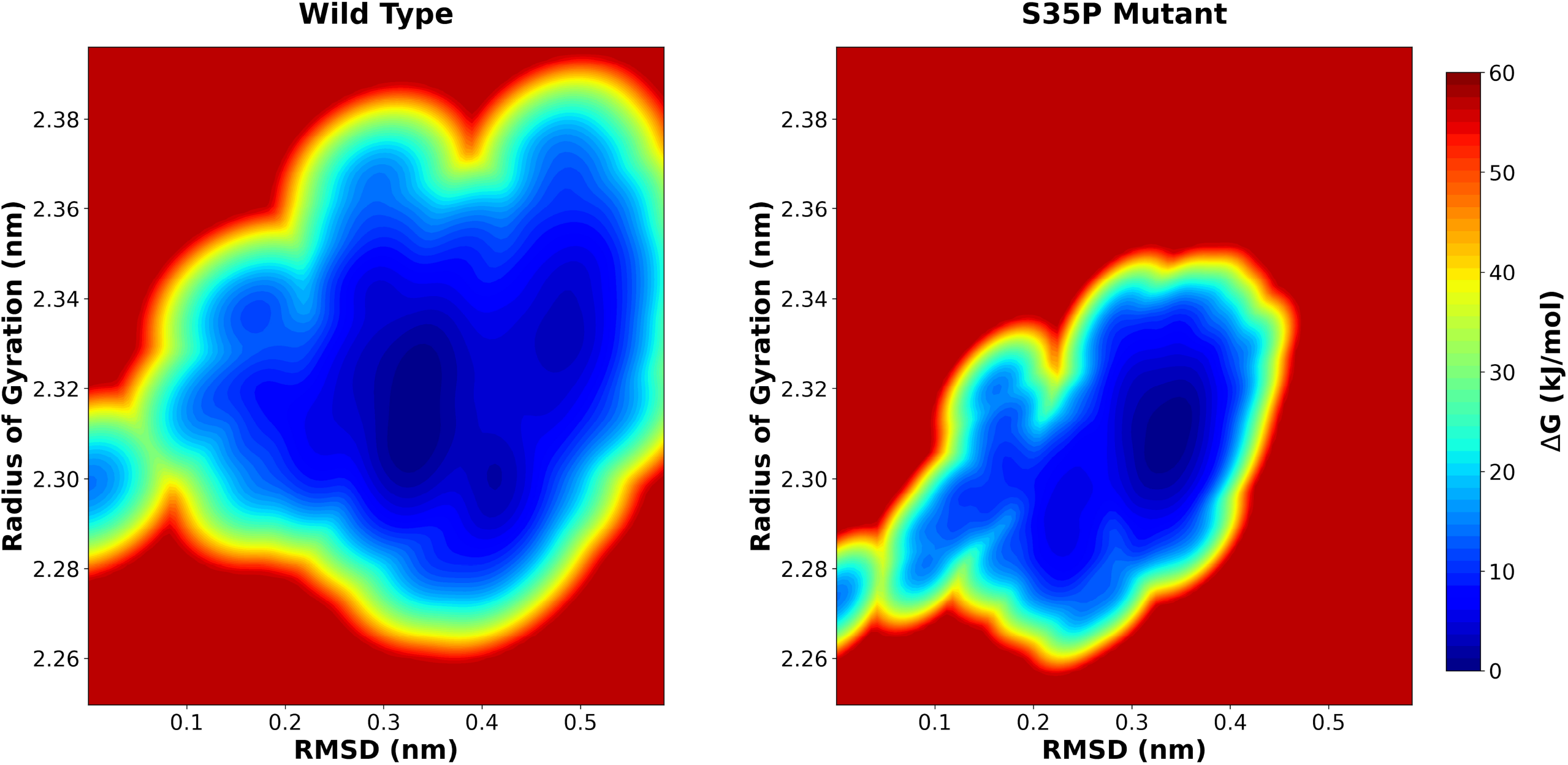
Free energy landscape (FEL) of WT and S35P MT-CYB derived from RMSD and Rg. Two-dimensional FEL maps for WT (left) and S35P mutant (right) projected onto RMSD (nm) and radius of gyration (Rg; nm). Color scale indicates relative Gibbs free energy (ΔG; kJ/mol), with lower values representing more frequently sampled, thermodynamically favorable conformations.

### 3.6. Thermodynamic Reorganization of the Heme bL Pocket

The energetic consequences of the structural and dynamic changes were quantified through MM-PBSA binding free energy decomposition (Fig. 7, Table S2). The S35P variant exhibited a decrease in van der Waals stabilization (−69.68 vs −62.43 kcal/mol), consistent with weakened local packing interactions. Most notably, the S35P variant was associated with a substantial reduction in electrostatic stabilization energy (ΔEEL), decreasing from +134.50 kcal/mol in the WT to +10.77 kcal/mol in the mutant (Table S2), in agreement with the disruption of the Ser35-mediated hydrogen-bonding network.

**Fig. 7.**
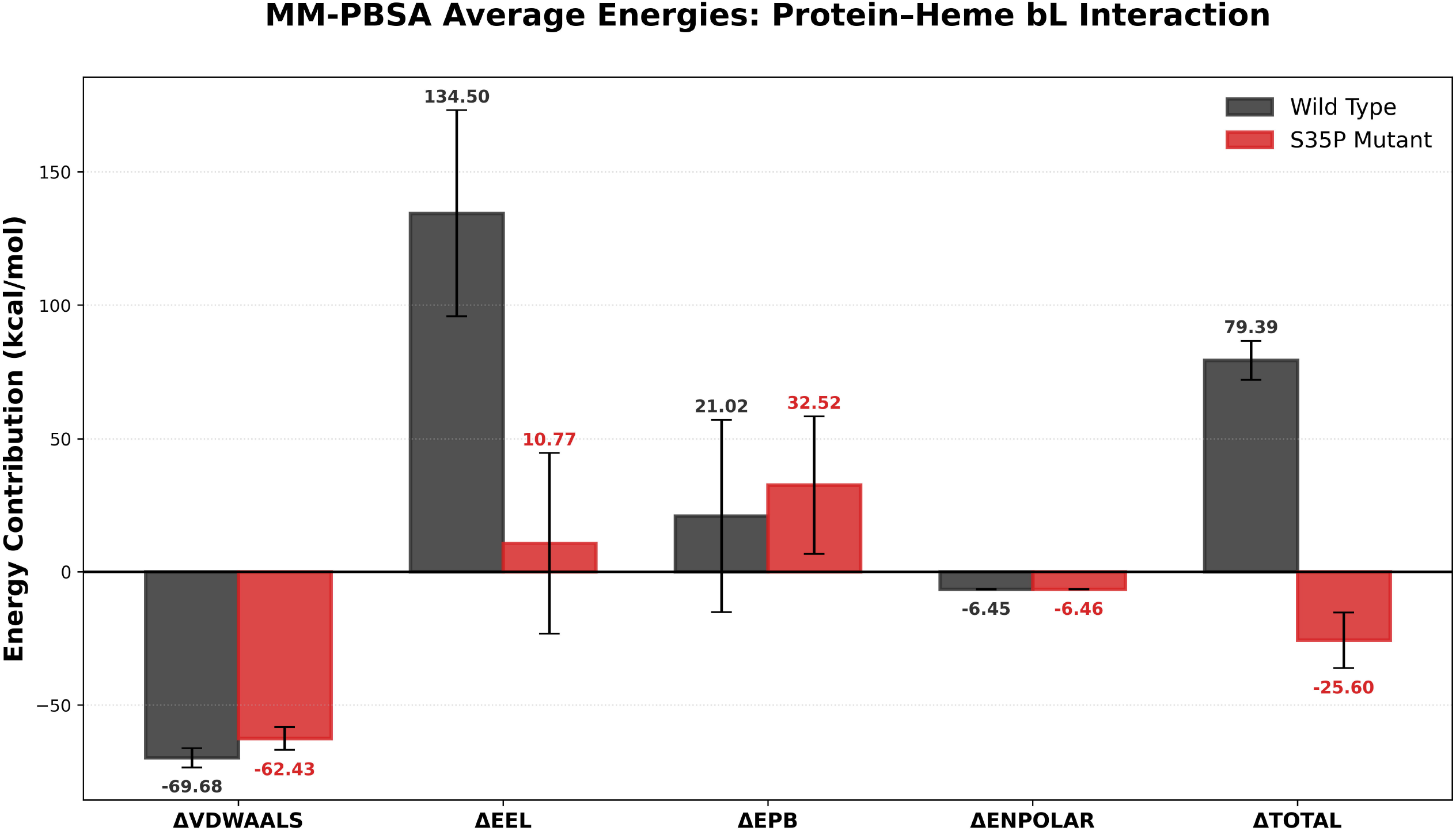
MM-PBSA decomposition of protein–heme bL interaction energies. Average energy contributions (mean ± SD) for van der Waals (ΔVDWAALS), electrostatic (ΔEEL), polar solvation (ΔEPB), non-polar solvation (ΔENPOLAR), and total binding energy (ΔTOTAL) for WT (gray) and S35P mutant (red) systems.

Despite the observed shift in total binding free energy (ΔTOTAL), the MM-PBSA estimates suggest a mutation-induced reorganization of the protein–heme interaction environment rather than enhanced stabilization. This interpretation is supported by the concomitant reduction in electrostatic contribution and the rightward displacement of the RDF coordination shell in the mutant system (Fig. S6), indicative of altered local packing around the heme bL cofactor.

### 3.7. Inter-Heme Geometric Perturbation Associated with the S35P Variant

Finally, we examined whether these microenvironmental alterations affect the inter-prosthetic geometry essential for electron transfer within Complex III. Distance monitoring between residue 35 and the heme bL cofactor revealed that the average separation remained geometrically comparable between WT and S35P systems throughout the simulation (Fig. 8A), suggesting that the mutation does not substantially alter the overall spatial positioning of this residue relative to the cofactor. In contrast, the inter-heme Fe(bL)–Fe(bH) distance displayed increased variability in the mutant, including transient shortening events not observed in the WT trajectory (Fig. 8B).

**Fig. 8.**
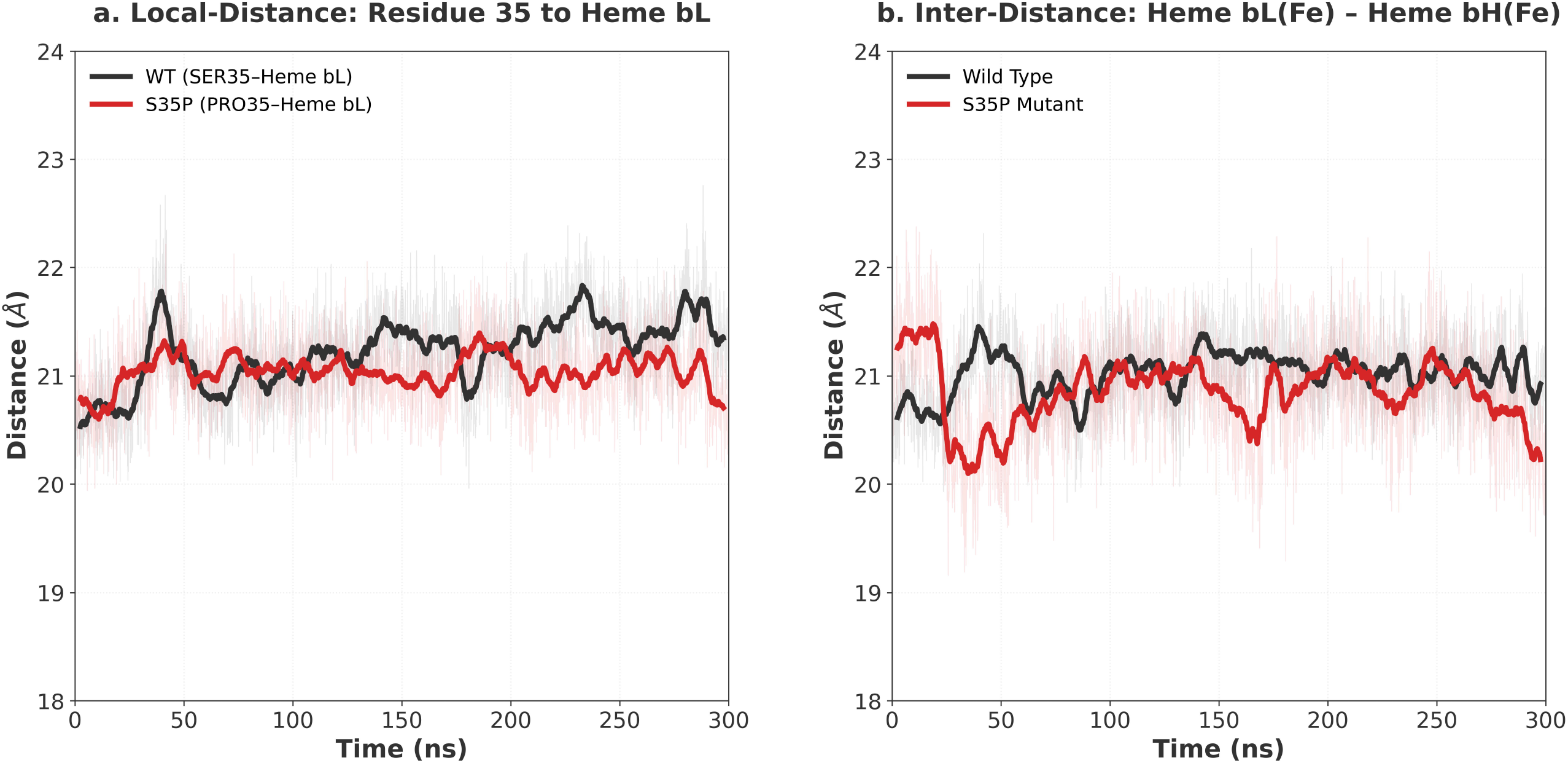
Local and inter-heme distance analysis in WT and S35P MT-CYB. (a) Time evolution of the distance between residue 35 and heme bL in WT (SER35) and S35P mutant (PRO35) systems. (b) Time-dependent Fe–Fe distance between heme bL and heme bH.

This geometric decoupling suggests that the localized helical perturbation at position 35 may influence the relative alignment of the redox-active centers. Such alterations in the inter-heme axis, in conjunction with reduced interaction stability and restricted conformational sampling, are consistent with a plausible structural mechanism that may contribute to reported Complex III–related phenotypes associated with the m.14849T>C variant.

## 4 DISCUSSION

Mitochondrial Complex III (cytochrome bc_1_ complex) plays a central role in oxidative phosphorylation by mediating electron transfer from ubiquinol to cytochrome c through the Q-cycle mechanism, a process that critically depends on the coordinated function of the cytochrome b subunit and its two redox-active prosthetic groups, heme bL and heme bH [4; 54; 55]. Variants affecting MT-CYB are known to impair Complex III activity and have been associated with septo-optic dysplasia, exercise intolerance, cardiomyopathy, and multisystem mitochondrial phenotypes [14; 16; 21]. However, for many reported missense substitutions including m.14849T>C (p.Ser35Pro) the molecular mechanisms linking genotype to dysfunction remain poorly defined [56-58].

In this study, atomistic MD simulations suggest that the S35P substitution does not induce global destabilization of Cytochrome b but instead alters the structural and energetic organization of the heme bL microenvironment. RMSD, Radius of gyration and SASA analyses (Fig. 2, Fig. S3, Fig. S4) indicate preservation of overall protein compactness, consistent with previous observations that pathogenic mitochondrial respiratory chain variants often impair function without causing large-scale unfolding of the affected subunit [59; 60]. Rather than promoting global instability, the S35P substitution appears to redistribute conformational dynamics within the transmembrane helical bundle.

Notably, RMSF and secondary-structure timeline analyses (Fig. 3, Fig. S4) revealed variant-associated flexibility changes in distal helices (residues 90–120) that form the structural scaffold surrounding the heme bL pocket. This observation is consistent with the concept that local perturbations within cytochrome b can propagate through the transmembrane helix network, influencing the relative positioning of the redox centers involved in electron transfer (Fig. 6A, Fig. 6B) [8; 61; 62]. Given that electron flow within Complex III proceeds sequentially between the bL and bH hemes during the Q-cycle [63; 64], even subtle alterations in the dynamic coupling between these regions may influence the efficiency of intra-protein electron tunneling.

Mechanistically, the primary molecular defect introduced by the S35P substitution appears to involve disruption of a heme-proximal hydrogen-bonding network. In the WT system, Ser35 participates in a stable polar interaction hub involving Asp228 and Asn32, which is markedly reduced in the mutant trajectory. MM/PBSA decomposition (Fig. 7, Table S2) further demonstrated a pronounced reduction in electrostatic interaction energy between the protein scaffold and the heme bL cofactor. This electrostatic reorganization was accompanied by a rightward shift in the RDF (Fig. S6), indicating loosening of local packing around the prosthetic group [65].

Importantly, distance monitoring revealed that while the mean separation between residue 35 and the heme bL cofactor remained comparable between WT and mutant systems, the inter-heme Fe(bL)–Fe(bH) distance (Fig. 8B) displayed increased variability in the S35P trajectory. Because the rate of electron transfer between redox centers is highly sensitive to both distance and relative orientation [66; 67], such mutation-associated geometric fluctuations may influence electronic coupling between the b-type hemes. These findings suggest that disruption of the Ser35-mediated interaction network may indirectly perturb the structural alignment of the inter-heme axis without necessitating large-scale conformational rearrangement.

Taken together, the present simulations support a model in which the m.14849T>C (p.Ser35Pro) substitution perturbs the dynamic organization of the heme bL pocket, leading to thermodynamic reorganization of the protein–cofactor interaction environment and altered inter-heme geometry. This localized microenvironmental effect is consistent with a plausible structural mechanism that may contribute to previously reported Complex III–related clinical phenotypes associated with MT-CYB variants [19; 68]. More broadly, these findings highlight how single-residue substitutions classified as variants of uncertain significance can exert measurable effects on cofactor-binding dynamics within mitochondrial respiratory chain proteins [69; 70].

## 5 CONCLUSION

Our study provides structural and energetic insights into the potential functional consequences of the MT-CYB m.14849T>C (p.Ser35Pro) variant, a missense substitution currently classified as a variant of uncertain significance. Our findings indicate that the S35P mutation does not induce global destabilization of the Cytochrome b subunit but instead promotes localized perturbation within the heme bL microenvironment. This variant-associated disruption of a heme-proximal hydrogen-bonding network leads to thermodynamic reorganization of protein–cofactor interactions and increased variability in inter-heme geometry.

These results suggest that even a single-residue substitution located within a heme-proximal transmembrane helix may influence the dynamic architecture of redox centers critical for electron transfer within Complex III. In this context, the observed microenvironmental perturbations provide a plausible structural mechanism that may contribute to previously reported multisystem mitochondrial phenotypes, including septo-optic dysplasia, cardiomyopathy, and exercise intolerance associated with the m.14849T>C variant.

Overall, this work highlights the importance of evaluating the structural and dynamic consequences of mitochondrial DNA variants classified as VUS, as localized perturbations in cofactor-binding environments may have functional implications without inducing global structural collapse. Future experimental studies will be necessary to determine whether these variant-associated microenvironmental perturbations contribute to the clinical phenotypes reported for this VUS through altered electron transfer dynamics in Complex III.

## Supplementary Information

Supplementary information containing additional analyses of structural dynamics, residue–heme interactions, and mutation-associated microenvironmental perturbations is available. The file includes 6 supplementary figures (Figs. S1–S6) and 2 supplementary tables (Tables S1–S2) supporting the molecular dynamics simulations and interaction analyses of the MT-CYB p.Ser35Pro variant.

## Acknowledgements

The authors acknowledge the computational resources provided by the Department of Biophysics at Erzincan Binali Yildirim University. Molecular dynamics simulations and data analyses were performed using local high-performance computing (HPC) facilities.

## Author Contributions

E.Y. conceived and designed the study, performed the molecular dynamics simulations, analyzed the data, and wrote the manuscript. A.Y.D. contributed to the clinical and genetic interpretation of the MT-CYB variant, analyzed the data, and critically revised the manuscript. S.D. contributed to study design, data interpretation, and critically revised the manuscript. All authors have read and approved the final manuscript.

## Data Availability

All data supporting the findings of this study, including molecular dynamics trajectories, analysis files and simulation movies are openly available in the Zenodo repository at https://doi.org/10.5281/zenodo.18775966 (accession number: 18775966).

## Declarations

### Ethics approval

Ethics approval statement is not applicable.

