## Supplementary material for "The Cytochrome b m.14849T>C (S35P) Variant Induces Structural and Dynamic Alterations in the Heme bL Microenvironment in Multisystem Disease": Supplementary_Information.docx

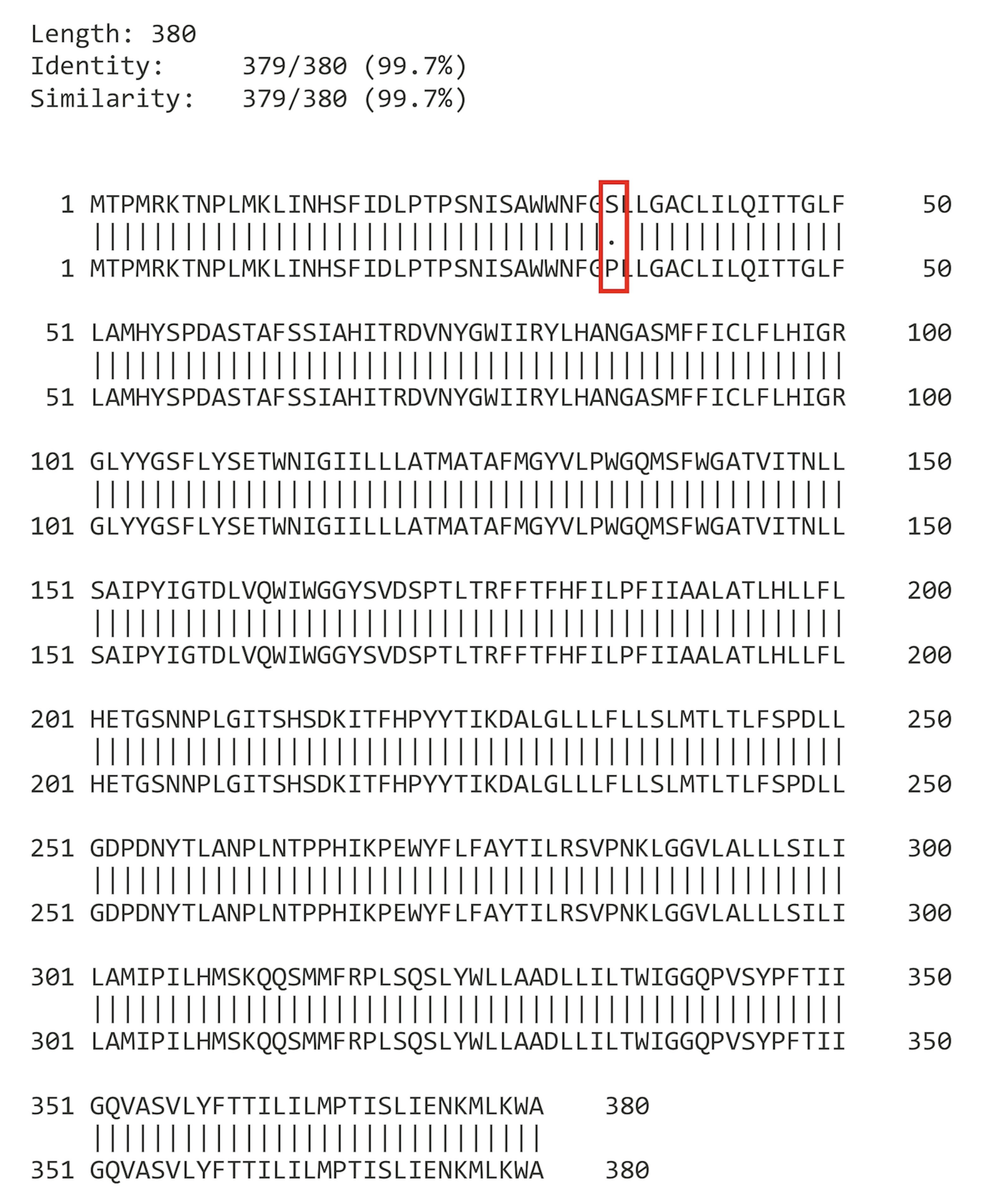

Figure S1. Pairwise sequence alignment of WT and S35P mutant human MT-CYB. The sequences show 99.7% identity (379/380), differing only at residue 35 where serine is replaced by proline (S35P), highlighted in red.

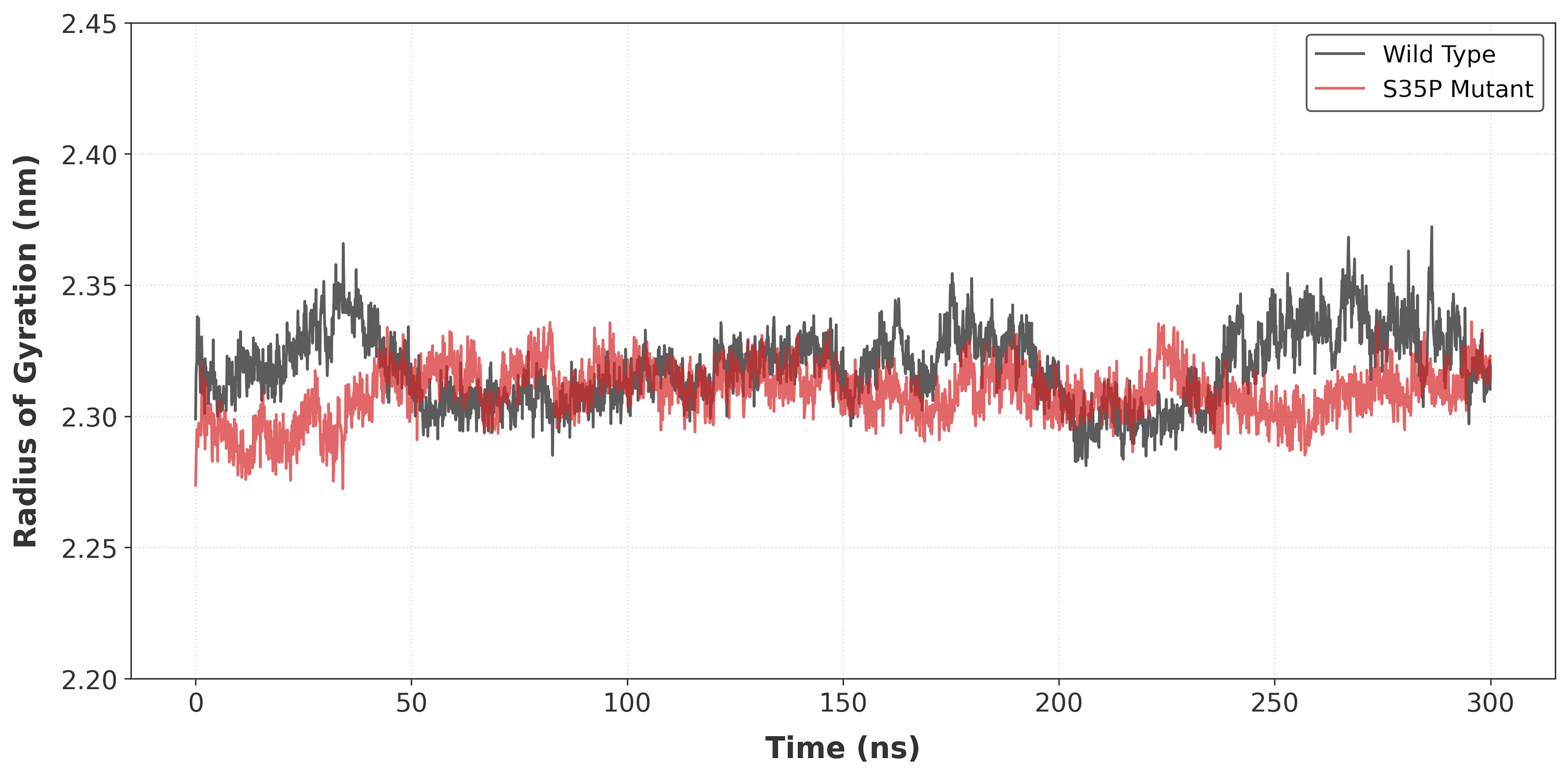

Figure S2. Radius of gyration (Rg) analysis of WT and S35P MT-CYB systems over 300 ns simulation time. Both systems retain comparable global compactness, indicating that mutation-induced effects are localized rather than globally destabilizing.

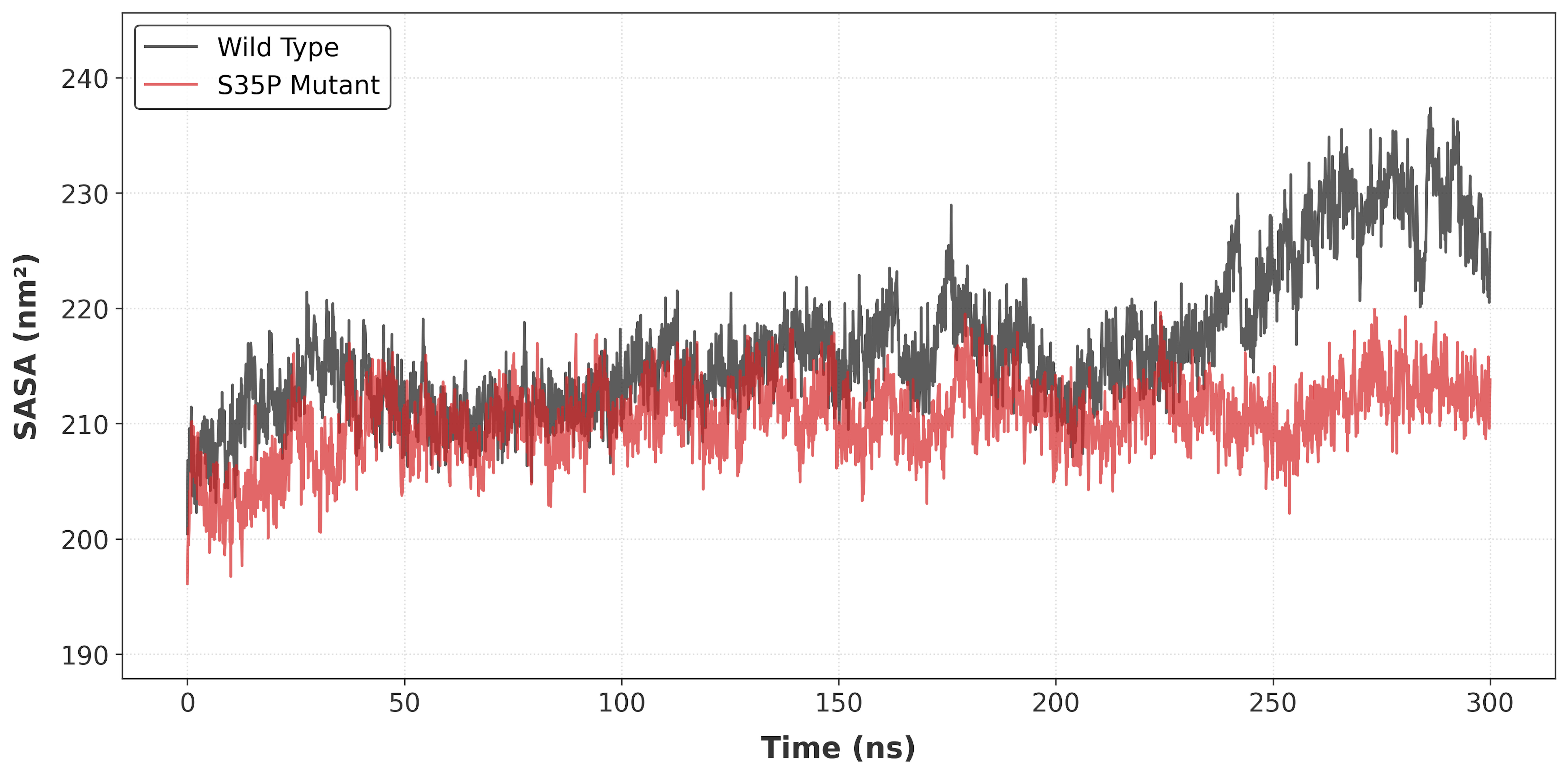

Figure S3. Solvent-accessible surface area (SASA) of WT and S35P MT-CYB systems over 300 ns simulation time. The S35P mutant exhibits reduced solvent exposure compared to WT, indicating mutation-induced restriction of local conformational flexibility.

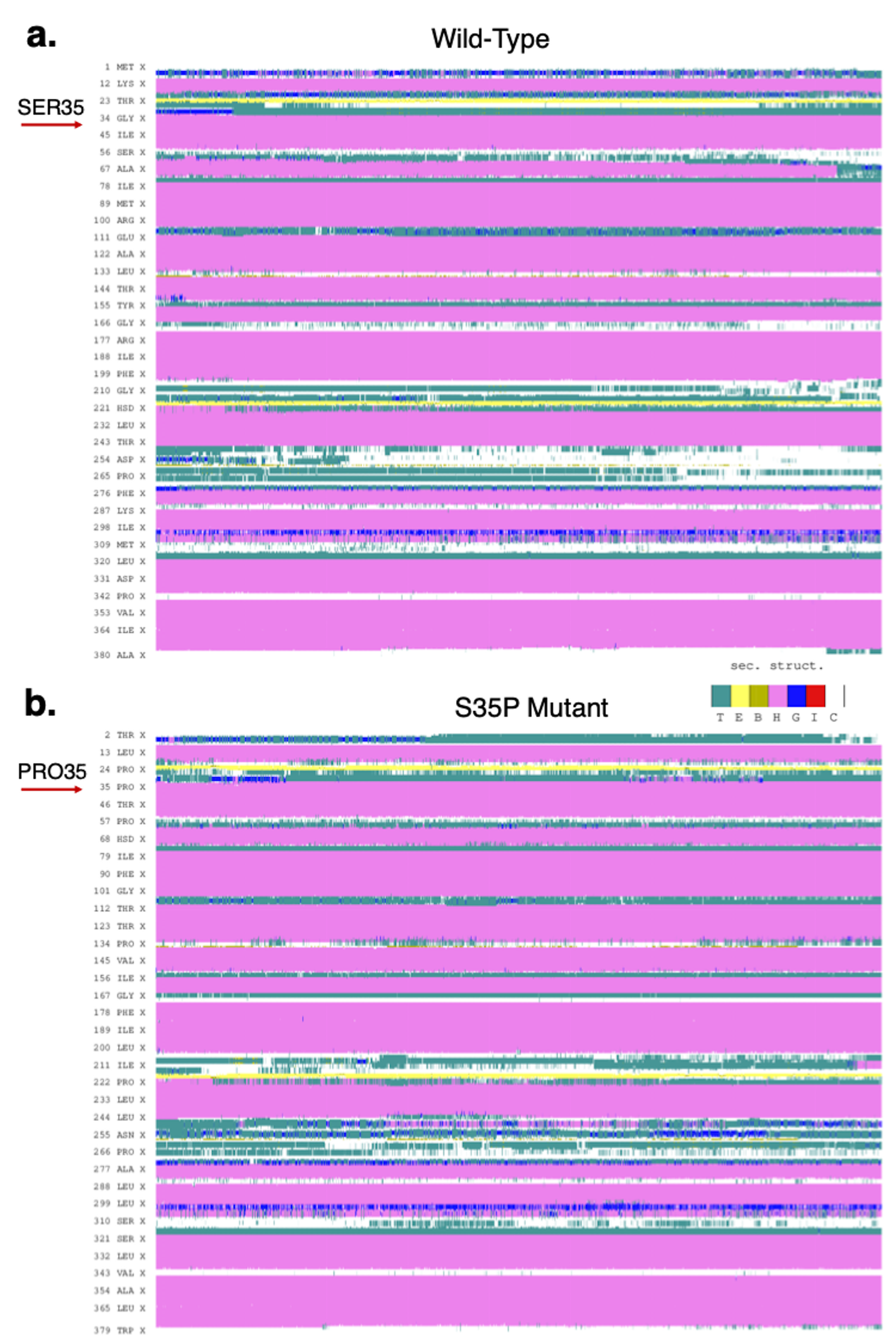

Figure S4. Secondary structure timeline analysis of WT and S35P MT-CYB systems over 300 ns simulation. The S35P mutant exhibits increased helix-to-coil transitions in regions proximal to the heme bL binding site, particularly within residues 90–120, indicating reduced local secondary structural stability.

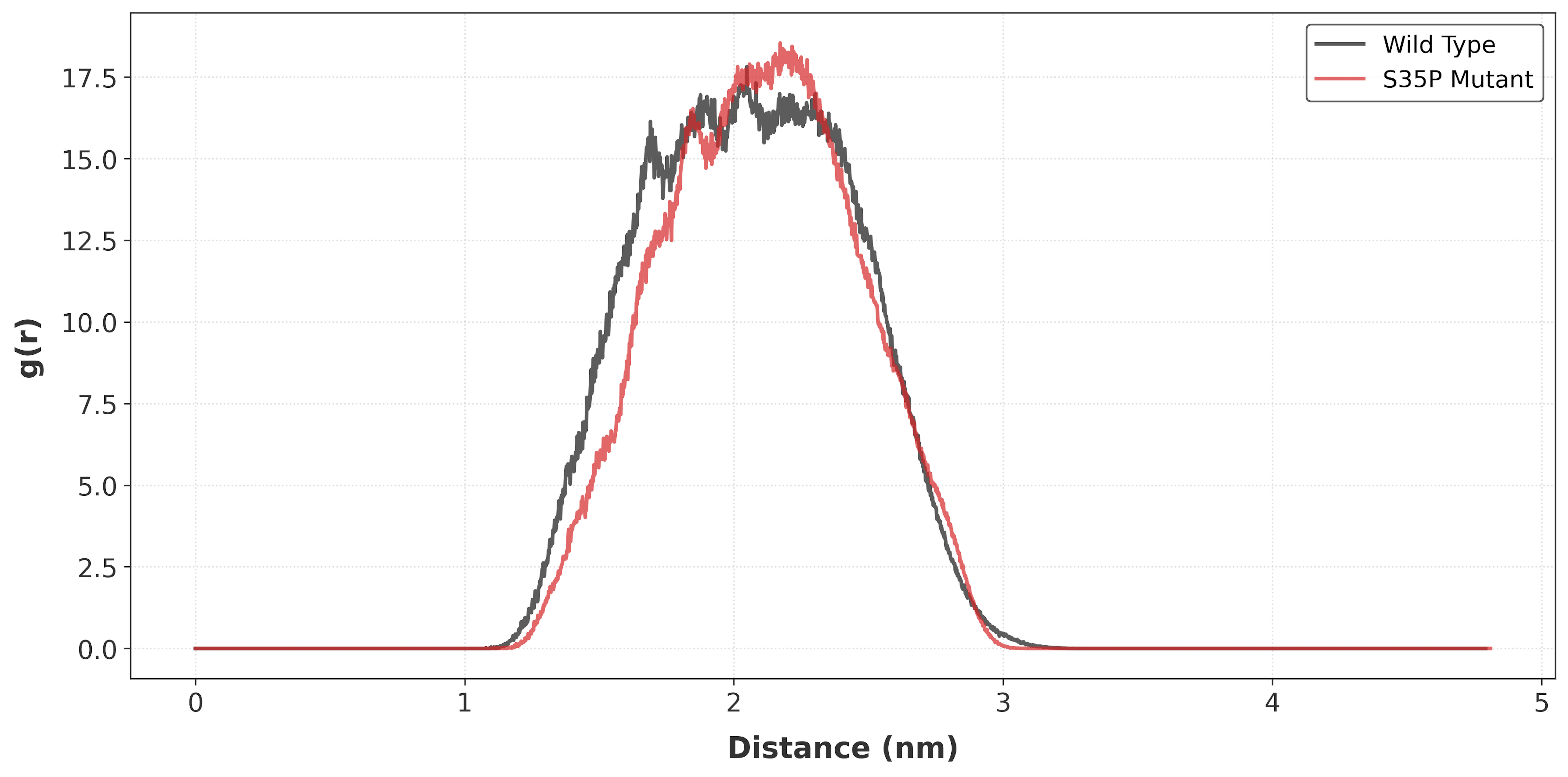

Figure S5. Radial distribution function (RDF) analysis between protein atoms and the heme bL cofactor in WT and S35P systems. The S35P mutation induces a shift in the primary coordination shell toward larger distances, suggesting reduced local packing density around the heme bL binding site.

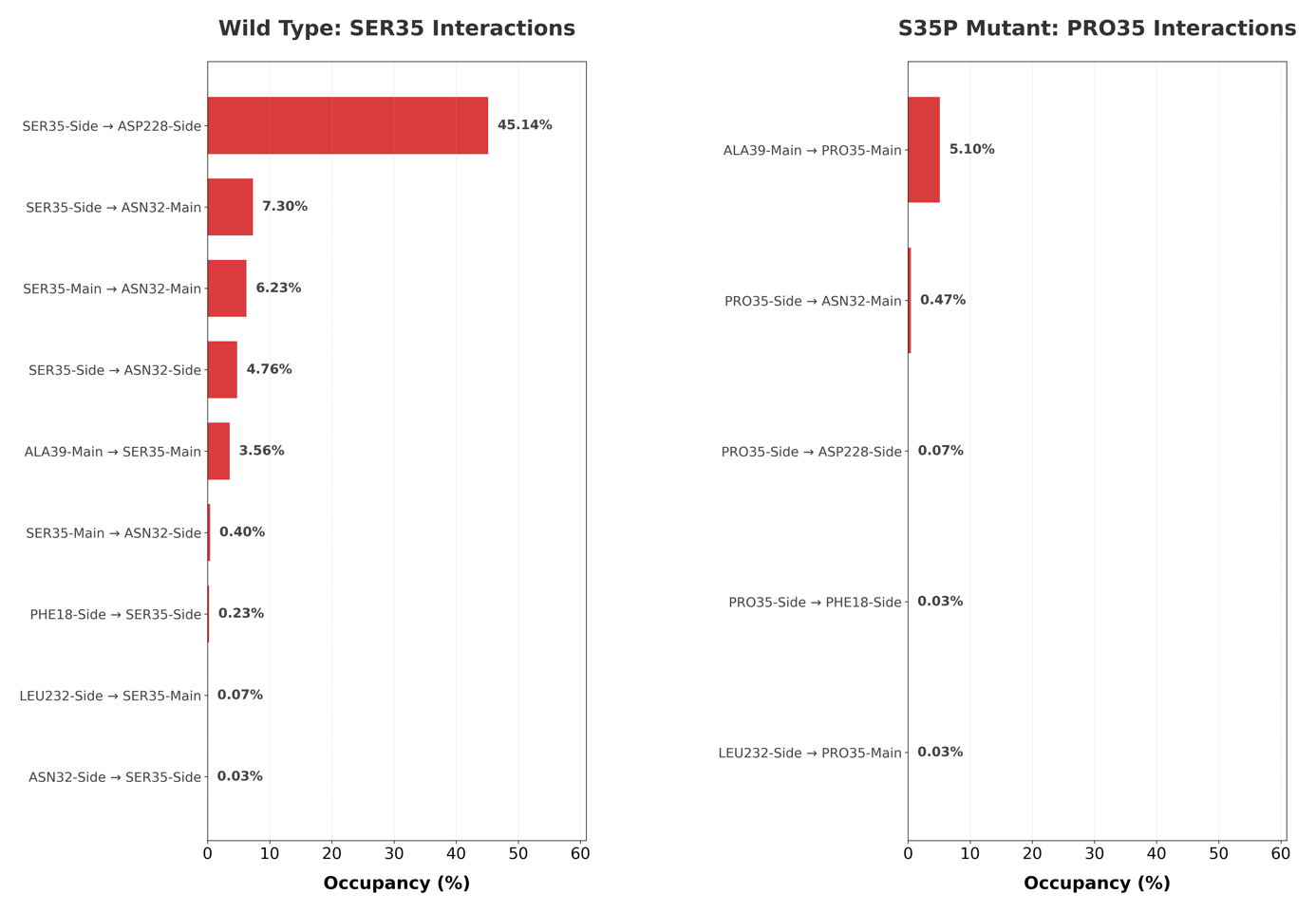

Figure S6. Hydrogen bond occupancy of residue 35 interaction partners in WT and S35P systems. WT SER35 forms a stable hydrogen bond network primarily with ASP228 and ASN32, whereas the S35P mutation markedly reduces interaction occupancy with surrounding residues, indicating disruption of local hydrogen bonding capacity.
