## Supplementary material for "The Cytochrome b m.14849T>C (S35P) Variant Induces Structural and Dynamic Alterations in the Heme bL Microenvironment in Multisystem Disease": Table_S1.docx

Table S1. Database Annotations of the MT-CYB m.14849T>C (p.Ser35Pro) Variant: Multisystem Mitochondrial Disease Phenotypes and Pathogenicity Classifications

| Locus | Nucleotide Position | Nucleotide  (AA Change) | Database | Clinical  Classification | Reported Disease / Phenotype | Main Clinical Features | References |
| --- | --- | --- | --- | --- | --- | --- | --- |
| MT-CYB | m.14849 | T>C (p.Ser35Pro) | ClinVar, MedGen | VUS | Mitochondrial Diseases; Exercise Intolerance; Cardiomyopathy; Septo-Optic Dysplasia | Developmental delay, hypotonia, ataxia, cerebellar hypoplasia, hypertrophic cardiomyopathy, retinitis pigmentosa, lactic acidosis, exercise intolerance, myopathy | [1], [2], [5], [8] |
|  |  |  | ClinGen | VUS | Mitochondrial Diseases | Developmental delay, lactic acidosis, ataxia, exercise intolerance, cardiomyopathy, retinitis pigmentosa, septo-optic dysplasia, cerebellar hypoplasia, mitochondrial myopathy | ClinGen CA120623, [5] |
|  |  |  | MITOMAP | Cfrm^*^ [VUS] | Exercise Intolerance / Septo-Optic Dysplasia | Developmental delay, hypotonia, ataxia, cerebellar hypoplasia, exercise intolerance, hypertrophic cardiomyopathy, retinitis pigmentosa, myopathy | [1], [2], [3], [4] |
|  |  |  | OMIM | Disease Associated  (Reported Variant) | Exercise Intolerance, Cardiomyopathy, Septo-Optic Dysplasia | Hypotonia, retarded language and motor development, dysdiadochokinesia, gait ataxia, microcephaly, cerebellar hypoplasia, exercise intolerance, cardiac hypertrophy, pigmentary retinopathy, optic nerve hypoplasia | OMIM #516020.0012, [6] |
|  |  |  | gnomAD | N/A | Not observed in population controls^**^ | — | [7] |

Abbreviations: AA, amino acid; Cfrm, Confirmed disease association (MITOMAP); VUS, Variant of Uncertain Significance; N/A, Not applicable.

^*^ VUS classification assigned by the ClinGen Mitochondrial Disease Nuclear and Mitochondrial Variant Curation Expert Panel (VCEP) per modified ACMG/AMP criteria (McCormick et al., 2020, DOI: 10.1002/humu.24107); curated June 2022.

^**^ Allele count = 0 in gnomAD v4.1 (61,883 mitochondrial sequences); absence in population databases supports pathogenicity. Data retrieved: February 2026.
