## Supplementary material for "The Cytochrome b m.14849T>C (S35P) Variant Induces Structural and Dynamic Alterations in the Heme bL Microenvironment in Multisystem Disease": Table_S2.docx

Table S2. MM-PBSA binding free energy decomposition (mean ± SD) for MT-CYB–heme bL interaction in WT and S35P systems.

| Energy Component (kcal/mol) | Wild Type (Mean ± SD) | S35P Mutant (Mean ± SD) |
| --- | --- | --- |
| ΔVDWAALS | -69.7 ± 3.6 | -62.4 ± 4.2 |
| ΔEEL | 134.5 ± 38.7 | 10.8 ± 33.9 |
| ΔEPB | 21.0 ± 36.0 | 32.5 ± 25.8 |
| ΔENPOLAR | -6.5 ± 0.1 | -6.5 ± 0.1 |
| ΔTOTAL | 79.4 ± 7.3 | -25.6 ± 10.4 |
